# First application of digital-PCR in oenology for the specific detection of intact cells of *Brettanomyces bruxellensis* in the winemaking process

**DOI:** 10.1101/2024.04.23.590681

**Authors:** Cécile Gruet, Jérémy Di Mattia, Magali Hiaumet, Dylan Pestel, Caroline Araiz, Sarah Saadi, Marie Ducousso, Olivier Courot

## Abstract

Wine is a complex matrix resulting from a fermentation process carried out by specific microbial communities. These communities can be in competition and the development of some microorganisms, as the yeast *Brettanomyces bruxellensis*, can impact the fermentation process and lead to organoleptic alterations of wine. To manage this risk, microbiological diagnostic methods as microscopic observations, qPCR or flow cytometry are already used in oenology, but remain either not specific enough, or tedious. In this context, IAGE (Ingénierie et Analyses en Génétique Environnementale) has developed the first digital-PCR system enabling the detection and quantification of *B. bruxellensis* during the whole winemaking process. Furthermore, wine DNA extraction was optimized to enable a representative and sensitive analysis of *B. bruxellensis* intact cells, as well as an easy-to-implement protocol to cope with the increasing number of samples to analyze. The IAGE workflow for *B. bruxellensis* quantification has been proven to be successful when analyzing naturally-contaminated samples during the different steps of the winemaking process and offers a robust method to oenologists for appropriate treatments and risk management in wine cellars.

**Highlights:**

- Development of a dPCR method led to a highly-specific analysis of *B. bruxellensis* intact cells in different steps of the winemaking process.
- DNA extraction method has been optimized to be robust across various types of wine with varying concentrations of inhibitors, as well as throughout different stages of the wine making process.
- The complete process was proven successful in analyzing a large number of naturally-contaminated samples, giving results in less than 48 hours.

## 1. Introduction

Microorganisms are essential for the winemaking process as they take part in alcoholic and malolactic fermentations. The wine microflora, evolving throughout the process, is mostly composed of bacteria and yeasts, which can be considered either as (i) beneficial for fermentation or (ii) wine contaminants, and are in constant interaction. On one hand, for example, *Saccharomyces cerevisiae* is known for its excellent oenological properties and is involved in alcoholic fermentation (Valero *et al*. 2007; Eliodório *et al*. 2019). On the other hand, contaminants microorganisms can cause wine spoilage by producing specific compounds, altering for example wine flavor, smell, or viscosity (Wedral *et al*. 2010).

Yeasts belonging to the species *Brettanomyces bruxellensis* (*B. bruxellensis*) are among these contaminants, causing major problems worldwide in wine production (Wedral *et al*. 2010). *B. bruxellensis*, also called *Dekkera bruxellensis* when being under teleomorphic (sexual) form (Van der Walt 1964), is ubiquitous and found throughout the winemaking process, from vineyard and grape vines to fermentation tanks and casks (Wedral *et al*. 2010; Curtin *et al*. 2015). The yeast, whose polyploidy status (50 % of available strains are triploid; Harrouard *et al*. 2023) is known to confer environmental stress resistance (Steensels, Gallone and Verstrepen 2021), also possesses several metabolic pathways, enabling it to thrive in diverse environments. These characteristics enable it to grow even in environments that have already undergone fermentation, with low-nutrient contents (Harrouard *et al*. 2023). Moreover, *B. bruxellensis* can tolerate high ethanol concentrations, low pH, and oxygen, making wine a conducive environment for its growth (Curtin *et al*. 2015), even though wines are known to be differently permissive to *B. bruxellensis* development depending on their chemical properties (alcohol concentration for example, Miranda *et al*. 2024). The pathway involved in wine alteration consists of the production of volatile phenol compounds (mostly 4-ethylphenol and 4-ethylguaiacol) responsible for altering wine’s organoleptic properties. These compounds are produced through the conversion of hydroxycinnamic acids, such as caffeic acid or p-courmaric, into vinylphenols (Zhang *et al*. 2021), and can result in “animal” aroma at high concentrations. In the first stages of vinification, *B. bruxellensis* is typically present at low concentration (competing with other yeasts responsible for alcoholic fermentation) but becomes more prevalent during malolactic fermentation (Renouf *et al*. 2006). It may increase to become the dominant yeast, thereby affecting the sensorial characteristics of wine, with a high risk of spoilage associated with a *Brettanomyces* concentration starting from 10^3^ CFU.mL^-1^ (Renouf and Lonvaud-Funel 2005). Considering all of this, *B. bruxellensis* characteristics necessitated investigation into its presence so risks can be anticipated, and appropriate treatments implemented to limit its proliferation.

*B. bruxellensis* physiology presents challenges for cultivation in traditional microbiological media due to its slow growth rate, viable-but-not culturable (VBNC) state, changing size and morphology; Serpaggi *et al*. 2012). The VBNC state of *B. bruxellensis* is primarily induced by environmental stresses, rendering the cells metabolically active but unculturable on standard media (Capozzi *et al*. 2016; Tubia *et al*. 2018). Consequently, these cells are undetectable on plates but remain capable of causing damage to wine, particularly following stress-inducing treatments (for example sulfur treatment). Furthermore, current microbiological analysis methods often involve growth on semi-selective media, leading to inaccurate results in terms of *Saccharomyces* or non-*Saccharomyces* yeasts counts after seven to ten days of incubation. Alternative methods, based on molecular biology, offer improved understanding of *Brettanomyces* spp. presence in the winemaking process (Pinto *et al*. 2020). For instance, flow cytometry coupled with fluorescence *in situ* hybridization (Röder *et al*. 2007), distinguishes viable and VBNC cells but presents challenges in terms of industrialization, while real-time PCR-based approaches (Cocolin *et al*. 2014, Tessoniere *et al*. 2009), requires an independent standard curve for each wine sample and may lack specificity, be time-consuming and be limited by PCR inhibitors present in wine. These limitations highlight the need for a fast, robust, and specific analytical process, capable of quantifying only intact cells to provide feedback, then allowing rapid treatments adjustment. Moreover, it must be possible to scale up such a process for industrial use. Digital-PCR, the latest generation of PCR technology, emerges as a promising candidate among available techniques. dPCR undergoes continuous development in human biology and is increasingly utilized in environmental biology (Condachou *et al*. 2024), offering reliable and absolute quantification, resilient to PCR-reaction inhibitors (Kuypers and Jerome 2017).

The objective of this study was to develop a Taqman-based tool for use in digital-PCR to specifically detect intact cells of *B. bruxellensis* for monitoring its prevalence in wine during the vinification process. To achieve this, we used available sequences of *B. bruxellensis* to design primers and probe for creating a detection system. The system was then assessed for its ability to specifically detect only this species and to cover the allelic diversity of the species simultaneously. Subsequently, we developed a wine pre-treatment for better DNA extraction and digital-PCR analysis process to provide a robust and reliable diagnosis of *B. bruxellensis* suitable for analyzing large volumes of samples, including all necessary quality controls for an industrialization of the process. We applied the developed method to approximatively 3000 wine samples from September 2023 to March 2024, highlighting the demand among oenologists’ for a rapid diagnostic method allowing early and appropriate treatments during the whole wine making process, and demonstrating prevalence of contaminants in French wine cellars.

## 2. Material and methods

### 2.1. Development of *in silico Brettanomyces bruxellensis* dPCR detection system

Sequence analysis and oligonucleotides designs were conducted using the Muscle algorithm for alignments (Edgar 2004) and Primer3 primer design program on Primer3plus website (Untergrasser *et al*. 2007), with specific parameters. *B. bruxellensis* specific primers and TaqMan probe were selected from the chromosome 5 sequence available on NCBI database, targeting a known differentiating zone allowing separation of *B. bruxellensis* from other species within genus. A consensus sequence was derived from alignment of 399 sequences from this zone. *In silico* specificity of primers and probe was evaluated using nucleotide BLAST (BLAST, http://blast.ncbi.nlm.nih.gov/Blast.cgi) against Genebank database.

### 2.2. Primer’s test and specificity validation on pure strain genomic DNA

To assess the ability of the designed primers and probe to specifically target a *B. bruxellensis* sequence, digital-PCR amplification was conducted on a QIAcuity Eight Plateform System (QIAGEN, Germany) following manufacturer’s instructions, using the QIAcuity Probe PCR Kit (Cat. No. 250103) and QIAcuity Nanoplate 26 K 24-wells (Cat. No. 250001).

The dPCR reaction mixture were prepared in a preplate as follows. For each reaction 10 µL of QIAcuity Probe PCR Kit, 1 µL of primers-probe mix, and 2 µL of *B. bruxellensis* CLIB316 genomic DNA were combined with H2O up to 40 µL final reaction volume. Reaction mixtures were transferred into a QIAcuity Nanoplate and the plate was loaded onto the QIAcuity Eight instrument, a fully automated system. The workflow included (i) priming and rolling step to generate and isolate the chamber partitions (26,000 partitions), (ii) amplification step with the following cycling protocol: 95°C for 2 min for the enzyme activation, followed by 40 cycles of 5 sec at 95°C for denaturation, and 30 sec of annealing/extension at 58°C, (iii) imaging step by reading fluorescence emission after excitation of the probe at the appropriate wavelength. Data were analyzed using the QIAcuity Software suite V1.2 and expressed as copies per microliter of reaction volume (40 µL).

Specificity of amplification was verified using microorganisms from the collection of the Institut de la Vigne et du Vin (ISVV, Villenave d’Ornon, France), provided by Microflora company (Villenave d’Ornon, France). Assays were done using 0.5 to 1 ng.µL^-1^ of gDNA from 16 different strains of *B. bruxellensis*, as well as 28 strains belonging to 18 other yeast species and 12 strains belonging to 9 bacteria species (Table 1). All strains were isolated from wine and wine processing environments, originating from diverse geographic locations, and varying phylogenetic distances from *B. bruxellensis*.

**Table 1.** List of yeast and bacteria species included in the detection system specificity test. Strains names were not given by ISVV and remained confidential. "+" indicates a dPCR amplification, "-" indicates no dPCR amplification.

| Species | Number of strains | dPCR amplification |
| --- | --- | --- |
| **Yeast** |  |  |
| *Brettanomyces bruxellensis* | 16 | + |
| *Candida boidinii* | 1 | - |
| *Candida cantarelli* | 2 | - |
| *Candida stellata* | 1 | - |
| *Debaryomyces hansenii* | 1 | - |
| *Hanseniaspora uvarum* | 2 | - |
| *Lachancea thermotolerans* | 2 | - |
| *Metschnikowia fructicola* | 1 | - |
| *Metschnikowia pulcherrima* | 2 | - |
| *Pichia kluyveri* | 1 | - |
| *Pichia membranifaciens* | 2 | - |
| *Saccharomyces cerevisiae* | 2 | - |
| *Shizosaccharomyces pombe* | 1 | - |
| *Starmerella bacillaris* | 2 | - |
| *Torulaspora delbrueckii* | 2 | - |
| *Trigonopsis variabilis* | 1 | - |
| *Zygosaccharomyces bailii* | 2 | - |
| *Zygosaccharomyces bisporus* | 2 | - |
| *Zygosaccharomyces rouxii* | 1 | - |
| **Bacteria** |  |  |
| *Acetobacter pasteurianus* | 1 | - |
| *Gluconoacetobacter liquefaciens* | 1 | - |
| *Gluconobacter cerinus* | 1 | - |
| *Lactobacillus brevis* | 2 | - |
| *Lactobacillus hilgardii* | 1 | - |
| *Lactobacillus plantarum* | 2 | - |
| *Oenococcus oeni* | 2 | - |
| *Pediococcus pentosaceus* | 1 | - |
| *Pediococcus parvulus* | 1 | - |

### 2.3. Method development for DNA extraction process

IAGE has developed a process to quantify only intact cells of *B. bruxellensis* enabling quantification of both viable cells (i.e. that can grow on Petri dishes) and VBNC cells. This process has been submitted for a FR patent (FR2401799). The protocol step is fully included into the pre-treatment before DNA extraction and is not detailed below. Tests for the efficacy of this process in red wine were performed as part of the patent (Table S1).

The initial DNA extraction process (Process A) was conducted using 100 µL of wine with the NucleoMag Tissue kit ® (Macherey Nagel, Oensingen, Switzerland) following manufacturer’s instructions, including an additional step for quantification of only intact cells before extraction.

The second DNA extraction process (Process B) was evaluated by incorporating additional steps to Process A to obtain a cleaner and more representative DNA extract. Two developments were implemented for this purpose. Firstly, various sample pre-treatments to remove unwanted residues were tested. A preliminary assay was conducted in duplicates using a characterized wine, for which *B. bruxellensis* concentration had already been measured. Four different washing solutions (more or less stringent with varying salt concentrations and with or without detergent) were tested before extraction with the NucleoMag Tissue kit®. A control with no treatment was also included. The composition of the solutions remains confidential and is proprietary to the IAGE company, hence detailed information is not provided. Once a solution was chosen, two different volumes of the solution (200 µL or 1.5 mL) were compared across 5 contrasting samples, without replicates, in comparison with no solution at all. Two of the samples were *B. bruxellensis*-negative (sample 1 and 2), two had low concentrations of *B. bruxellensis* (sample 3 and 4), and one was a finished wine in a bottle, therefore containing fewer residues, supplemented with a high and known concentration of yeast in liquid-culture (sample 5). Secondly, the sampling volume of wine was increased to be more representative of the total sample (200 mL are received for analysis). To achieve this, five different sampling volumes of wine were compared (100 µL, 500 µL, 1 mL, 1.5 mL, or 2 mL), with or without treatment with the previous chosen solution, in duplicates. The chosen wine sample for this test was the finished wine in a bottle. These preliminary assays helped choose the optimal combination of sampling volume and treatment.

With this chosen sample pre-treatment parameters, a new assay was conducted. First, fourteen *B. bruxellensis*-negative wine samples were chosen among several wine samples that had followed extraction Process B and analysis by dPCR in our lab. Then, the experiment proceeded as follows: a liquid-culture of *B. bruxellensis* CLIB 316 at a known concentration was used to prepare four different dilutions in Phosphate Buffer Saline solution (PBS 1X), to create a concentration range between 0 and approximatively 120 copies. µL^-1^ in dPCR reaction (3/4 dilution, 1/2 dilution, 1/4 dilution, and 1/8 dilution). Then, for each dilution, three different samples were each supplemented with 20 µL of the dilution, in triplicates, giving nine replicates (3 samples x 3 replicates) per tested *B. bruxellensis* dilution. All replicates were tested in two conditions: with or without the washing steps. In addition to these samples, one *B. bruxellensis*-negative wine sample was used as a negative control, supplemented with 20 µL of PBS 1X in triplicates, and one sample was also used as a positive control, supplemented with 20 µL of the pure, non-diluted, *B. bruxellensis* liquid culture (i.e. routine positive control). All prepared samples were then extracted and analyzed by dPCR to obtain quantification results. The R 4.3.3 software was used for statistical analysis, with a chosen p-value *p* < 0.05. Data and residuals were checked for normal distribution (Shapiro-Wilk test) and homoscedasticity (Levene test). *B. bruxellensis* concentrations were analyzed using a one-way ANOVA approach and Tukey’s HSD test to compare washed / non-washed conditions. Then an ANOVA of a nested-model followed by a Tukey’s HSD test was used to compare (i) washed and non-washed condition for each sample within each dilution category (considering variability between replicates of each sample) and (ii) the global effect of the washing steps within each dilution category (considering variability between samples inside each category).

### 2.4. Inhibition test

Digital-PCR step was also optimized to increase the volume of total DNA extract analyzed to improve representativity of the measure. As inhibition can be observed in PCR amplification due to wine components, inhibition tests were conducted to test PCR inhibitors hypothetical presence on four wine DNA extracts extracted with Process A and containing already characterized *B. bruxellensis* concentrations. These samples varied in terms of quantity and origin, sourced from different wine terroirs to ensure representativeness of sample diversity. dPCR was performed as described in section 2.2 but a range of DNA extract from 1 to 10 µL was added to the reaction mix and inhibition curves were produced.

### 2.5. Method characterization and quality control setup

The smallest copy number of the targeted sequence detectable in a reliable manner per dPCR well (hereafter called absolute limit of detection) was determined. One sample was selected and analyzed thrice to calculate the mean *B. bruxellensis* concentration. The sample was then diluted to achieve final concentrations corresponding to 20, 10 and 5 DNA copies per dPCR reaction (copies deposited in plate wells). Each dilution was measured in three replicates on the same dPCR run.

Then, the smallest number of *B. bruxellensis* to be detectable starting from raw samples (200 mL), in a reliable manner, was determined (hereafter called global limit of detection). To do so, from a *B. bruxellensis*-negative wine sample, three sub-samples of 200 mL each were supplemented with a *B. bruxellensis* liquid culture at a final concentration of 1.58 x 10^4^ copies. mL^-1^, three others at a final concentration of 7.90 x 10^3^ copies. mL^-1^ and the last three at a final concentration of 4.00 x 10^3^ copies. mL^-1^, hence giving three independent replicates for each chosen concentration. Each replicate was then extracted in triplicates and analyzed by dPCR. This finally resulted in nine replicates of analysis for each level of *B. bruxellensis* concentration in a raw sample of 200 mL.

To ensure reliable analysis for wine cellars and normalize *B. bruxellensis* quantification, proper quality controls were added to check process’s smooth running. For the extraction workflow, a known sample of stable wine from a bottle, not containing any *B. bruxellensis,* was supplemented with a known concentration of yeast in liquid-culture and added to each extraction run. This extracted spiked sample was added to each dPCR run, along with another control consisting of a *B. bruxellensis* DNA extract at a known concentration. Both control concentrations were monitored to ensure consistency. Additionally, a negative control was included in each extraction and analyzed by dPCR to check for any contamination during the extraction process.

### 2.6. Comparison of *Brettanomyces bruxellensis* dPCR and conventional-plate culture method

Forty-two samples underwent both culture-dependent and culture-independent methods for the detection and quantification of the yeast. On one hand, the Institut Coopératif du Vin (ICV, Lattes, France) analyzed each sample following their IGA method (Spoilage germ index) for the detection of colonies of *Brettanomyces*-like yeast (expressed in CFU.mL^-1^), on Petri dish (non-*Saccharomyces* medium). On the other hand, we performed DNA extraction with Process A (see details in section 2.3) from 100 µL of the same wine samples. Extracted DNA samples were analyzed by dPCR following the optimized protocol described above and results were expressed in DNA copy. mL^-1^. Using the IGA method as a reference, dPCR results were compared to the colony-counting method. To verify results, plates were sent to IAGE and Petri-dish colonies were analyzed by dPCR to confirm or refute *B. bruxellensis* identification. This secondary experiment was to check colonies identity, so three identical colonies of each morphotype were directly resuspended into 50 µL H2O and this suspension was used as a template in the reaction mix without prior DNA extraction.

### 2.7. *Brettanomyces bruxellensis* dPCR analysis in naturally contaminated samples

Before testing naturally contaminated samples, the ability to run a great number of samples at the same time was assessed. DNA extraction, after pre-treatment process, was performed on an automated extraction system allowing simultaneous extraction of 96 samples. dPCR was both done on 26 K 24-wells or 8.5 K 96-wells plate (Cat. No. 250021), in order to compare the ability to accurately quantify serial dilutions of *B. bruxellensis* even on a 96-wells plate, where less dPCR reactions are performed (8,500 instead of 26,000 per well on a 24-wells plate). Obtained results for 24-wells plate and 96-wells plate, only for washed samples, were compared with a Spearman correlation test to determine the significance of the correlation between values and the correlation factor.

A total of 2988 wine samples, over seven months (from September 2023 to March 2024), were then analyzed following the developed workflow (Process B) to test our *B. bruxellensis* analysis by dPCR.

## 3. Results and discussion

### 3.1. Development of a digital-PCR detection system for a specific *Brettanomyces bruxellensis* quantification

Creating a suitable detection system for *B. bruxellensis* in digital-PCR is based on designing two primers and a TaqMan probe. The three oligonucleotides were scrutinized against NCBI databases and showed no significant homology with any “wine-organisms” other than *B. bruxellensis*. The forward primer and probe exhibited 100 % coverage and identity exclusively with *B. bruxellensis* (95 % coverage for forward primer corresponded to other eukaryotes organisms as marine organisms or insects), while the reverse primer showed 100 % coverage only with a clam species. The use of a probe increases theoretical specificity as the fluorescence signal depends on probe hybridization on target DNA. A previous study reported the development of qPCR primers for the specific detection of *B. bruxellensis* (Phister and Mills 2003, Tessonière et al. 2009), later used in other studies (Willenburg and Divol 2012), however some non-specific amplification of *Saccharomyces* species was sometimes reported (Tessonière et al. 2009). This highlighted the need to test the experimental specificity of our dPCR system.

Designed oligonucleotides enabled amplification of a 234 bp fragment and produced a fluorescence signal when used on pure strain *B. bruxellensis* CLIB 316, confirming its targeting ability. Specificity was confirmed using gDNA from yeast and bacteria isolated from wine and wine process environment, from contrasted geographic origins. On the sixteen *B. bruxellensis* strains targeted, dPCR assay gave a positive signal for all of them (Table 1). The ability to detect a wide variety of strains confirms that the system targets the all-species diversity. This is crucial because despite *B. bruxellensis* intraspecific diversity, partly explained by strain isolation origin (wine, beer, kombucha, etc.; Avramova et al. 2018) due to genetic adaptation to processes, wine isolates are distributed into different genetic groups (Harrouard et al. 2023). CLIB 316 was, on the other hand, initially isolated from beer, demonstrating that our detection system can be valuable for assessing fermentation process beyond wine. No amplification was detected for the other eighteen yeast species and nine bacteria species tested (Table 1), thus, confirming the method’s specificity. Indeed, among the tested yeast species some are more phylogenetically distant as *Zygosaccharomyces* spp. while others are genetically close to *B. bruxellensis*, as *Pichia membranaefaciens* (Cai *et al*. 1996), which is not targeted by our dPCR system (Table 1).

### 3.2. Characterization of inhibitors effect on dPCR amplification

While dPCR is more resilient to inhibitors than previous PCR technologies, inhibition can still happen to some extent. Inhibition tests were conducted on samples previously extracted with Process A. The goal was to assess the hypothetical presence of residual inhibitors in DNA extract after our washing process, and in a second time, to enhance the representativity of the measure by increasing the DNA extract volume in the dPCR reaction mix, while not inducing a loss in quantification. The original volume in the reaction mix was 2 µL, and higher volumes were tested (4, 6, 8 and 10 µL). For all samples, the quantification from 4 µL of DNA extract was equal to the quantification obtained with 2

µL (data not shown). For 3 of the 4 contrasted samples, the optimal volume seemed to be 8 µL as a decrease of concentration was observed for some of them at 10 µL (Figure 1A). However, for one sample, concentration instantly lowered when the volume increased above 4 µL (Figure 1A). Inhibitors concentration differs depending on wine type or wine process (Wilson 1997; Zoecklein *et al*. 1999). This may lead to varying residues concentration in DNA extract, with a possible lower efficacy of the developed washing treatment depending on the wine type. Our goal was to be able to offer a reliable analysis for all wine types even though DNA extract are not perfectly clean. Hence, in order to choose the optimal DNA volume, the sample with the highest inhibition was used for a new assay with 1, 2, 3, 4 or 5 µL of DNA extract. This time, a real lower quantification was observed only at 5 µL (Figure 1B). However, a decrease in fluorescence intensity between 3 and 4 µL conditions was observed during results analysis (approximatively 75 RFU (Relative Fluorescens Unit) compared to approximatively 90 RFU for the cluster of positive dPCR partitions; data not shown), indicating a minor inhibition. It was then decided to make a compromise and set the volume at 3 µL to avoid any kind of inhibition. Therefore, all subsequent analyses were carried out with 3 µL of DNA extract in a 40 µL final volume of reaction mix, and DNA extraction pre-treatment tests were validated with this dPCR protocol.

**Figure 1.**
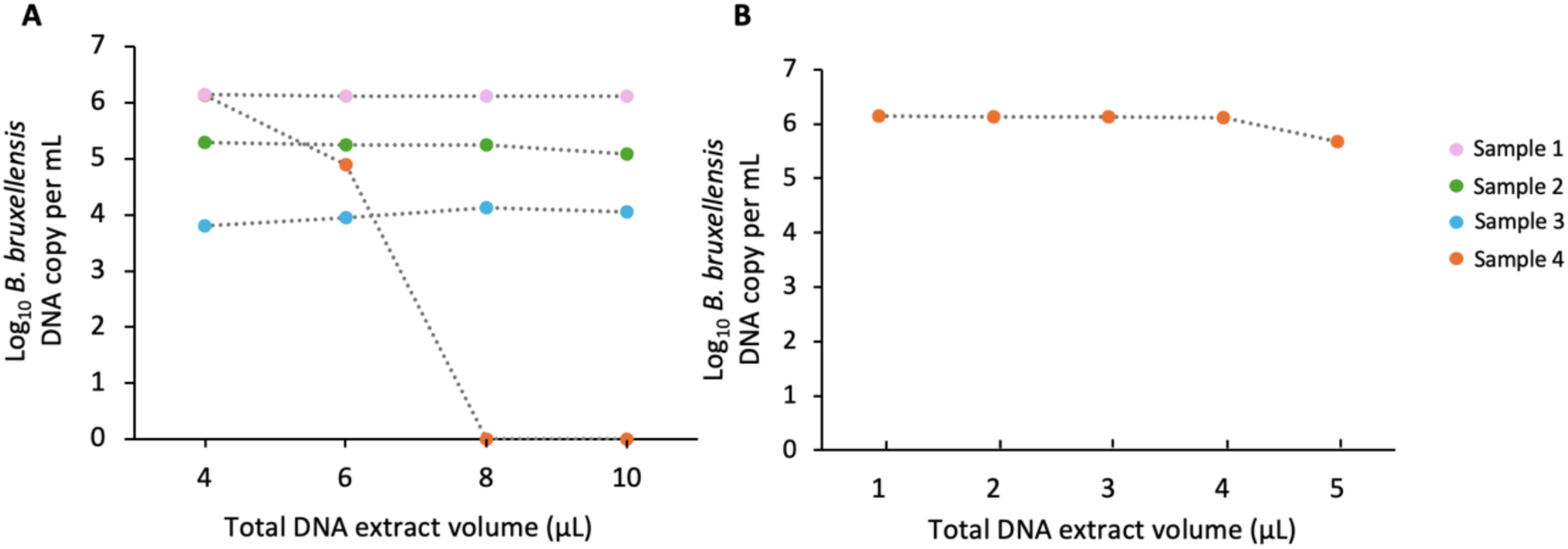
*Brettanomyces bruxellensis* dPCR concentration according to DNA extract volume in the reaction mix. (A) The different colors correspond to different samples. The original tested volume being 2 µL, other tested volumes were 4, 6, 8 and 10 µL. (B) Orange sample was tested again with 1, 2, 3, 4 or 5 µL of DNA extract.

### 3.3. Development of reliable extraction process for a relevant analysis of *Brettanomyces bruxellensis* intact cells in wine

As dPCR requires DNA extraction, which is not necessary with conventional plate-cultures, it must be optimized to avoid any loss of signal due to extensive manipulations. The analytical process was developed to obtain a robust method for wine diagnosis, even when a large number of samples are analyzed simultaneously. Indeed, when analyzing environmental samples, the quality of DNA extraction must be considered. Previous methods have been tested for DNA extraction from wine to lower inhibitors concentrations in DNA extracts as it is a complex matrix containing undesired residues (Tessonnière *et al*. 2009; Jara *et al*. 2008).

Here, the objective was to create a reliable and rapid workflow, ready for industrialization. Our process involves a pre-treatment step to quantify only intact cells in different kind of environmental samples. This step was tested on wine before any further experiment as part of patent FR2401799 submission. Table S1 is provided as an example of result, showing that, in a *B. bruxellensis*-positive wine sample supplemented with *B. bruxellensis* gDNA, the yeast DNA copy concentration is lower when samples are treated following our process than when no treatment is applied. Another method, based on qPCR quantification of mRNA was developed by Willenburg and Divol (2012), to quantify viable cells and VBNC cells. This method was proven effective in accurately detecting VBNC state cells. However, mRNA extraction remains expensive and difficult to industrialize, in comparison to our process which was fully included in the tests described below.

Preliminary assays were conducted to determinate a representative pre-treatment that would eliminate PCR-inhibitors. We initially extracted of 100 µL of wine (Process A) to correlate with what is achieved in conventional plate assays in the litterature (Tubia *et al*. 2018). Different pellet washing solutions after centrifugation of 100 µL of wine were tested. Solution C, the more stringent one (containing higher concentration of salts and detergent), was the only solution to allow a higher quantification of *B. bruxellensis* than the control sample where no treatment was applied, increasing quantification from 2.57 x 10^3^ to 6.19 x 10^3^ DNA copies.mL^-1^ (Figure S1 A), hence this solution was chosen for subsequent analysis. The volume of washing solution can have a great impact, therefore, several volumes were tested on five contrasted samples, positive samples containing either low or high concentration of *B. bruxellensis* or negative samples for *B. bruxellensis*. The washing steps had no impact on the two negative samples 1 and 2 and did not increase nor lessened the quantity of *B. bruxellensis* in the bottle wine sample (sample 5; Figure S1 B). This result confirms that no loss of quantification is induced by the washing process on samples that are already rather clean. However, for samples 3 and 4, washing the pellet with 1.5 mL of solution C increased quantification in comparison to not washing the pellet (Figure S1 B). Quantification was slightly higher for sample 4 with 200 µL of solution C than with 1.5 mL (1.11 x 10^3^ vs 6.37 x 10^2^ copies.mL^-1^), but it was decided to keep 1.5 mL as 200 µL might not be enough for other types of wine (i.e. sample 3; Figure S1 B).

The second strategy to improve DNA extraction was the increase the sampling wine volume, from 0.1 mL to 2 mL, as this has an impact on *B. bruxellensis* detection and quantification variability (Willenburg and Divol, 2012). We obtained rather close concentrations ranging between 5.11 x 10^3^ and 8.13 x 10^3^ copies.mL^-1^, regardless of the sampling volume (Figure S1 C). Considering it was tested on a finished wine in bottle, no significant differences between washed and non-washed samples were expected. Additionally, it is forecasted that a small sampling volume mostly affects low concentrated *B. bruxellensis* samples. Indeed, a low sample volume decreases the representativeness of the sampling, increasing the likelihood of missing a few yeast cells, and consequently leading to greater variability in the measurement. Hence, the impact of sampling volume is not observed in this highly-concentrated sample. Therefore, it was decided to stick to the 1.5 mL sampling for the industrialization of the method for two reasons. Firstly, it enhances the representativeness of the measurement, multiplying the volume tested by fifteen, if compared to the volume usually plated (100 µL) in conventional plate assays (Phister and Mills, 2003; Tubia *et al*. 2018). Secondly, risks of contamination are lower than with 2 mL because microtubes are not filled to the top, which allows a better practicality.

To confirm the preliminary assays a second test was conducted, this time including fourteen samples tested in triplicates with the validated washing treatment or no washing. Among these fourteen samples, two were controls, a positive one (spiked with a characterized *B. bruxellensis* liquid culture) and a negative one. Additionally, three were supplemented with 3/4 dilution of the pure culture, three with 1/2 dilution, three with 1/4 dilution and three with 1/8 dilution. Supplementing three different wines with the same concentration of *B. bruxellensis* allowed to consider contrasting types of wine in the assay, as residues, inhibitors, and polyphenols concentrations can vary between wines of different origins (Makris *et al*. 2006). *B. bruxellensis* DNA copy number was indeed correlated with the supplemented yeast quantity (Figure S2A). Firstly, considering all results, *B. bruxellensis* dPCR quantification was significantly higher for washed compared to non-washed samples (Figure 2). Then, considering each dilution category separately, washed samples showed a significantly higher quantification than non-washed samples for pure culture (category A), 3/4 dilution (category B), 1/2 dilution (category C) and 1/4 dilution (category D) (Figure 2), meaning at the smallest dilution (1/8, category E), no significant effect of the washing treatment was observed. However, washing does not induce a loss of quantification, as a lower concentration was observed only for sample 11 when washed, but a higher variability is observed between replicates of this sample (Figure 2). Pre-treatment was significantly effective for sample 1 and 4, as well as for the positive control (Figure 2). Our process seems to be suitable for samples highly concentrated in *B. bruxellensis*, while not inducing loss for low-concentrated samples, enabling the analysis of a wide range of wines. This also demonstrates all phenolic compounds produced by the yeast (Cibrario *et al*. 2020), which are major PCR inhibitors, are mostly washed away with the pre-treatment. Therefore, combining this pre-treatment with an optimized dPCR sampling volume, together with dPCR being an inhibitor resilient method by nature, the process enables an accurate quantification. All these improvements in wine samples pre-treatment led to the adoption of Process B for DNA extraction, currently used in the laboratory.

**Figure 2.**
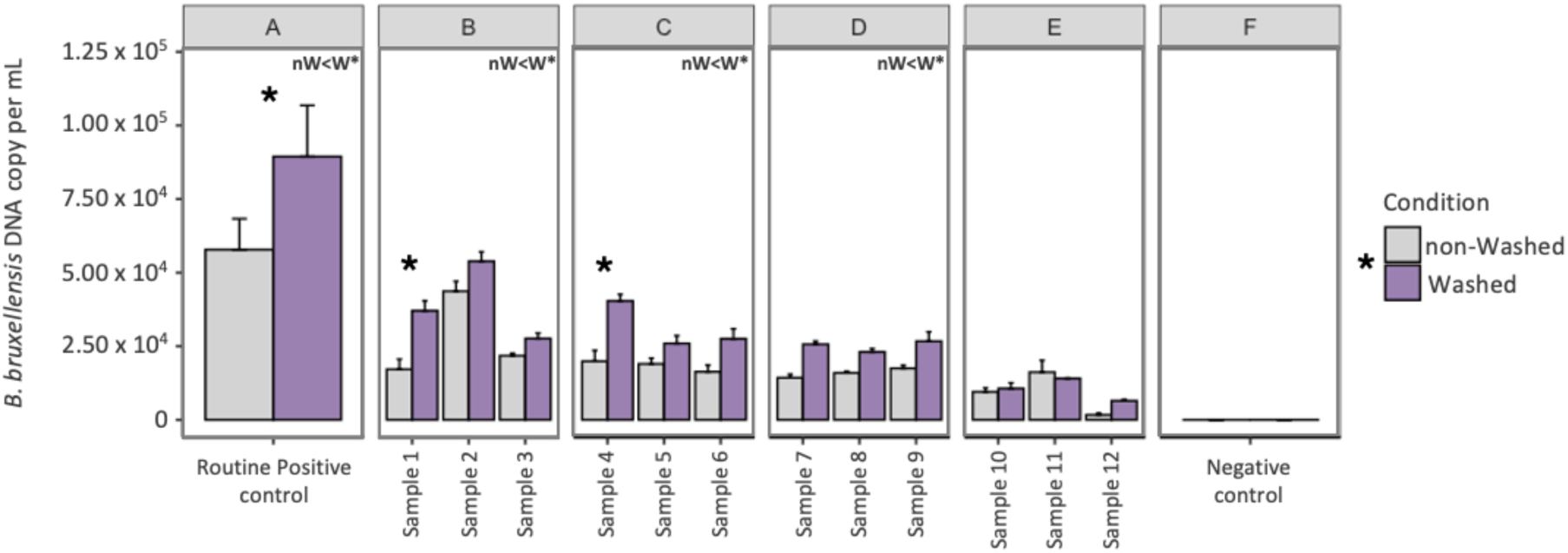
Process B confirming experiment for the quantification of *B. bruxellensis* in wine. For each sample, dPCR concentration means are represented and standard errors are calculated from n=3 for A and E, n=6 for B, C and D. A: Routine positive control corresponds to a *B. bruxellensis* wine negative sample inoculated in triplicates with 20 µL of *B. bruxellensis* liquid culture at a known concentration. B: samples 1, 2 and 3 were inoculated with 20 µL of 3/4 dilution of pure *B. bruxellensis* liquid culture. C: samples 4,5 and 6 with 1/2 dilution. D: samples 7, 8 and 9 with 1/4 dilution. E: samples 10, 11 and 12 with 1/8 dilution. Samples were either washed according to our Process B (purple) or non-washed (grey). “*” indicates statistic significant difference set at *p* <0.05. ‘nW<W’ indicates that levels were significantly higher for washed samples compared to non-washed samples.

### 3.4. Absolute and global limits of detection for *Brettanomyces bruxellensis*

On the three replicates conducted to establish the absolute detection limit, all of them exhibited a fluorescence signal at the three tested dilutions, *i.e.* 20, 10 and 5 copies per dPCR reaction. Since the validation of a limit of detection is based on qualitative criterions, positive result was obtained for all three replicates, the smallest copy number of the targeted sequence that the dPCR method can detect was validated at 5 copies per dPCR reaction.

With extraction Process B, we were able to determine the global limit of detection, *i.e.* the smallest number of *B. bruxellensis* to be detectable in raw samples (with maximal size of starting sample being 200 mL), in a reliable manner. On the nine replicates tested for each of the three concentrations, all of them showed a fluorescence signal at the three tested dilutions, *i.e.* 1.58 x 10^4^ copies. mL^-1^, 7.90 x 10^3^ copies. mL^-1^ and 4.00 x 10^3^ copies. mL^-1^. The global detection limit was set at 4.00 x 10^3^ copies. mL^-1^. In previous studies, detection limits were determined around 1 to 5 CFU / mL (Phister and Mills 2003; Tessonnière *et al*. 2009; Tofalo *et al*. 2012; Willenburg and Divol 2012). However, it remains challenging to compare detection limits with previous studies since one CFU corresponds to several cells and then several DNA copies of the targeted zone in this assay. Regarding flow cytometry, it has been shown that its detection limit remains higher than with conventional-plate culture or PCR-based approach (Bouix *et al*. 2022).

### 3.5. Comparison between *Brettanomyces bruxellensis* dPCR analysis and conventional-culture and plate counting methods

*B. bruxellensis* analysis is usually conducted using plate assays and counting CFU on a yeast-specific medium. To confront digital-PCR method to conventional analyses of wine contaminants, both culture-dependent (plate assay) and culture-independent (dPCR) methods were performed. A total of forty-two red wines were subjected to plating on a non-*Saccharomyces* medium (IGA method, developed by the Institut Coopératif du Vin) and to DNA extraction followed by dPCR analysis. IGA method was considered as the reference method and dPCR results were interpreted accordingly. Considering qualitative data (detection/no detection), about 33 % of samples gave identical results between both methods (Table 2). *B. bruxellensis* was detected on eighteen plates (considered *B. bruxellensis*-positive) and the other twenty-four plates showed no yeast growth (considered *B. bruxellensis*-negative; Table 2). It is important to note that results from the IGA method are expressed in term of “*Brettanomyces*-type yeast” and the medium is not selective for *B. bruxellensis* species only. Among the eighteen positive plate samples, only 8 were determined positive by dPCR and 10 negatives. Among the twenty-four negative plate samples, *B. bruxellensis* dPCR amplification was observed for six of them (Table 2) and concentration ranged between 7.40 x 10^1^ and 1.40 x 10^4^ DNA copies. mL^-1^ (Table S2). Quantification data were generally not correlated between plate counts and dPCR (Table S2).

**Table 2.**
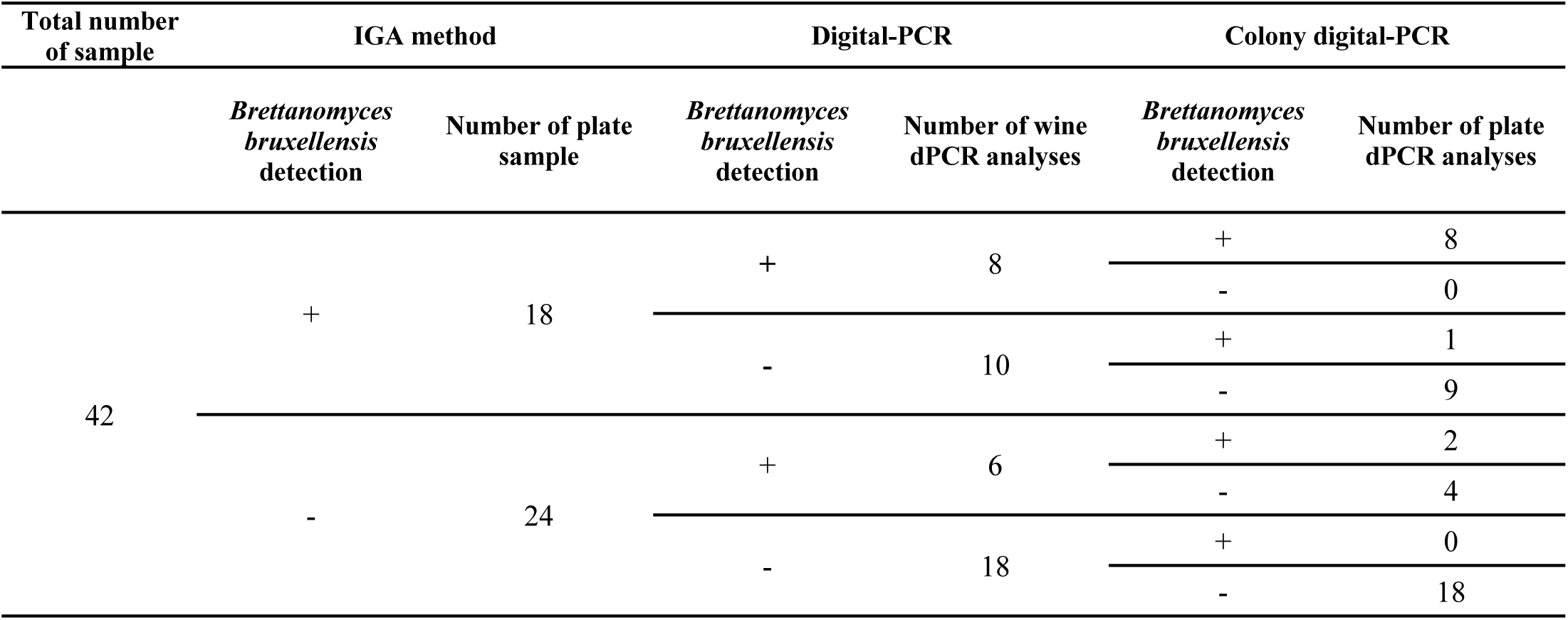
Second comparison of *Brettanomyces bruxellensis* detection in red wines according to two different methods: IGA plate method and dPCR and validation with colony dPCR. A total of 42 red wine samples were selected and analyzed. IGA method was developed by the Institut Coopératif du Vin and consists in conventional plate assay. dPCR analyses were made on DNA extract from 100 µL of the same wine samples. IGA method was considered as the reference and dPCR was compared to it. Colonies on Petri dish were retrieved and analyzed by dPCR. "+" indicates presence of > 1 CFU on Petri dish or > 1 positive dPCR partition. "-" indicates no CFU on plates or no dPCR amplification. Quantification details are given in Supplementary Table S2.

After the seven days of incubation, all plates were subjected to colony-dPCR analysis. This experiment confirmed nine samples out of the eighteen *B. bruxellensis-positive* plates indeed harbored *B. bruxellensis* colonies (Table 2), but the rest most probably corresponded to other yeast species since non-*Saccharomyces* medium is not a selective medium. On the six *B. bruxellensis*-negative plates, determined positive by dPCR, two were truly corresponding to *B. bruxellensis* (colony-dPCR results; Table 2). Moreover, on the eighteen samples both determined as negative by IGA and dPCR methods, all of them were true-negative (Table 2). Only one sample determined positive by plate assays but negative by dPCR gave an amplification with the colony-dPCR (Sample 30; Table S2). This contrasted result can be explained by the low *B. bruxellensis* concentration in the sample, probably being under the detection limit in dPCR, but getting positive because amplification is easier from plate colonies.

Differences between both methods can partly be explained by the volume of wine analyzed. Indeed, sampling 0.1 mL from a total volume of 200 mL might lack representativeness, leading to increased variability in results. Several previous studies compared plate count and PCR amplification of *B. bruxellensis.* Although a study reported that a good correlation between plates and qPCR can be observed (Phister and Mills, 2003), depending on wine type, no correlation is observed (Tofalo *et al*. 2012; Willenburg and Divol, 2012). The hypothesis of a small sampling volume is supported by data from Willenburg and Divol (2012), where no *Brettanomyces* is detected on plates from 0.1 mL of wine, and detection is seen with qPCR analysis from 2 mL of wine. Moreover, due to VBNC state of *B*. *bruxellensis*, plate counts can give false-negative results (Agnolucci *et al*. 2009), whereas dPCR can detect cells in this state. It has been observed that a long winemaking process with sulfiting can lead to development of smaller microorganisms that forms microcolonies not visible in only seven days of incubation (Millet and Lonvaud-Funel, 2000). Therefore, the fact that we were able to quantify *B. bruxellensis* in samples where plates showed no yeast growth, might be explained by either variability due to small volumes or VBNC state. When colony growth was observed but no dPCR amplification happened, this was most probably due to the non-specific medium which allows other yeast species, that are neither *Brettanomyces bruxellensis*, nor *Saccharomyces* spp., to grow. This hypothesis was partly confirmed by colony dPCR.

A correlation between DNA copies and CFU per mL is often calculated, creating a standard curve to convert qPCR results in CFU.mL^-1^ (Tessonnière *et al*. 2009, Willenburg and Divol 2012). This standard curve approach aims to align result with the unit of the conventional plate-culture method, which remains the gold standard worldwide. However, we argue this is not mandatory since this type of standard curves are calculated from only one strain, while it is well known that, because of different polyploidy levels, different *B. bruxellensis* strains will have more or less DNA copy number (Harrouard *et al*. 2023). Therefore, relying on one strain to calculate a correlation may not accurately represent the variability existing between samples. Having a diagnostic based on accurate quantification data avoids this bias.

Out of all 42 samples tested, 24 % would have been declared positive with the IGA method and treated against *Brettanomyces*, despite apparently being indeed negative. Although the IGA method can lead to an overestimation and unnecessary treatments in a context of treatment reducing policies, it remains a reliable estimation method. It is more qualitative than quantitative but has been extensively used in the past and possesses a good reference database and interpretation keys. However, dPCR allows a clear and faster (approximatively 24 to 48 hours compared to 7 days) identification of wine contaminants with less doubts, besides detection of only intact cells thanks to our DNA extraction process. This faster application has allowed oenologist to implement dPCR analyzes as soon as the grapes are picked in order to prevent wine contamination by *B. bruxellensis*.

### 3.6. Industrialization of the process

To evaluate the potential industrialization of our method, we first needed to determine if a large number of samples could be processed simultaneously in a reliable manner. DNA extraction can be performed using an automated extraction system capable to extract 32 or 96 samples. Previous investigations into the homogeneity of extraction between the two systems showed no differences (data not shown). Additionally, the dPCR analysis can be conducted either on a 24-wells plate (performing 26,000 reactions per well) or 96-wells plate (performing 8,500 reactions per well). We investigated whether similar quantities *B. bruxellensis* could be retrieved regardless of the number of reactions. To do this, the same fourteen samples tested in section 2.3. were analyzed on a 96-wells plate to compare to the 24-wells plate results. As for the 24-wells plate results, *B. bruxellensis* DNA copy number was correlated with the supplemented yeast liquid culture quantity (Figure S2A). Moreover, DNA copy number results from both type of plates were significantly correlated, indicating that we were able to retrieve approximatively the same tendency of concentration, regardless of the number of dPCR reaction (Spearman’s correlation factor = 0.94, p > 0.05; Figure S2C). However, we observed an average loss of 31 % with the 96-well plate in comparison to the 24-wells plate. This suggests that the method is quite resilient to different types of material, even when many samples need to be analyzed simultaneously. However, subsequent analyses of naturally contaminated samples were done on 24-wells plate, to ensure a workflow that can be industrialized depending on the number of samples received each day, while giving a reliable and rapid result, avoiding any bias.

### 3.7. Analysis of naturally-contaminated samples

The complete workflow, from sample pre-treatment to dPCR analysis, was executed on nearly 3000 samples on a seven-month period. During the peak demand period, 237 samples were analyzed (week 43), over 5 days, averaging forty-seven samples per day (Table 3). Furthermore, the industrialization of the process enabled reliable results for 130 samples from a single wine cellar in just 48 hours. The substantial number of samples in September, October and November can be partially attributed to the grape-harvest season and the initial stages of winemaking process, as *B. bruxellensis* is known to be present on grapes (Curtin *et al*. 2015), and contamination could occur through these first steps. On a total of 2988 wine samples, 60 % were *B. bruxellensis*-negative and 40 % positive (Table 3). The percentage of negative samples increased over time, month after month (Table 3). This could be related to treatments applied to wine to reduce *B. bruxellensis* contaminations, advice was indeed provided by oenologists based on our quantification data after each analysis. However, since this is a blindfolded monitoring, further information about samples is not available. Among samples received between September 2023 and March 2024, some of them were white and rosé wine. These samples were treated as red wines, although additional positive controls for these specific samples were added to each run of analysis. In the same manner as for red wine, white or rosé wine were supplemented with liquid culture of the yeast and included in the process. This enabled the detection and quantification of *B. bruxellensis* in other types of wines, retrieving the expected quantity of yeast after spiking. The necessity of using internal controls was already described in the literature (Tessonnière *et al*. 2009), since it is impossible to know in advance if samples contain a high concentration of inhibitors or not. Furthermore, our process was successful in analyzing other winemaking process steps, to extract DNA from sample with a higher residue content, like wine must at the end of grape-harvest season. Regarding these samples containing high concentrations of inhibitors, we were able to apply our washing process to 100 µL sampling volume and to control DNA extraction’s smooth running (data not shown).

**Table 3.**
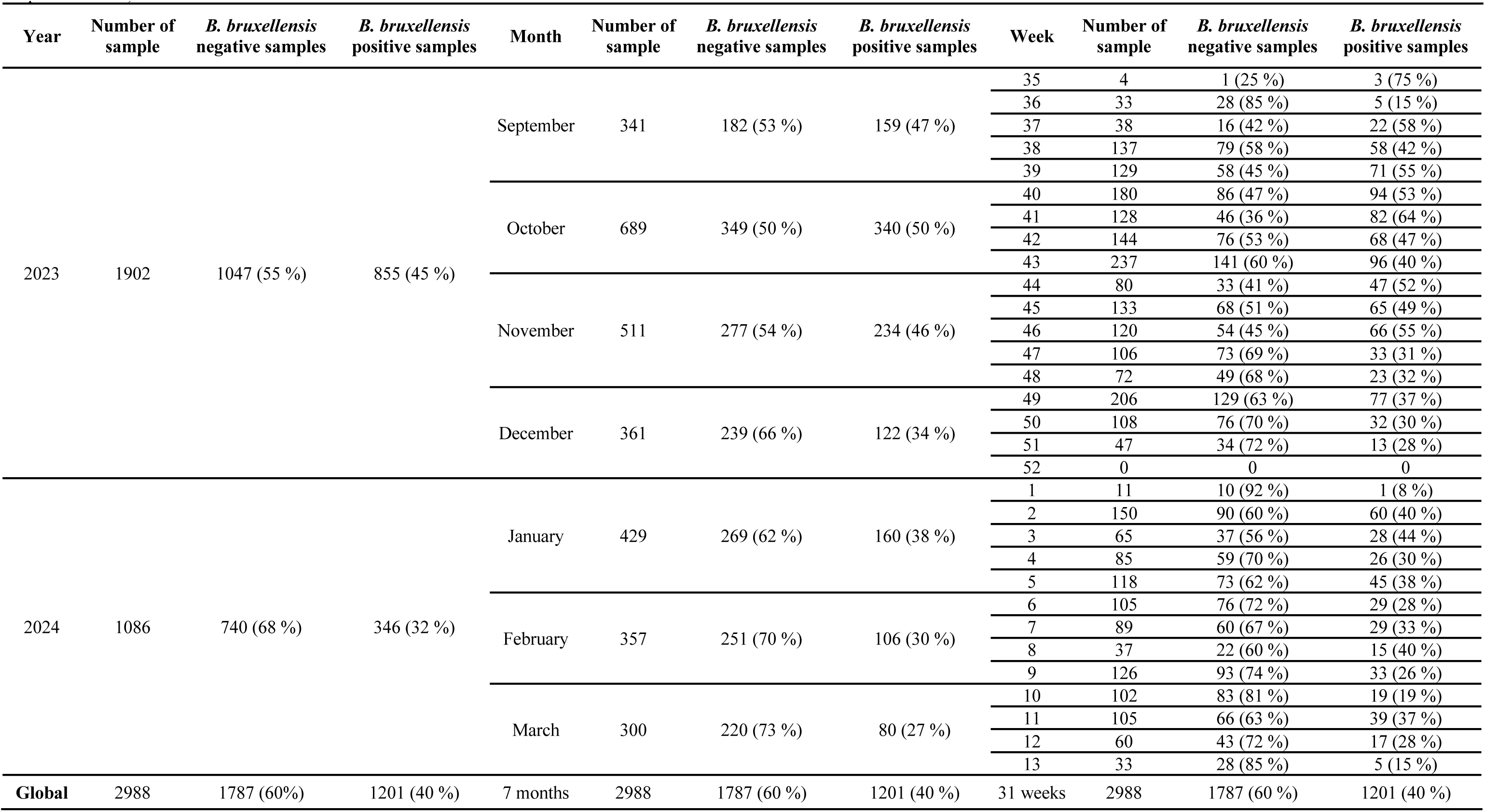
dPCR *Brettanomyces bruxellensis* quantification in 2988 naturally-contaminated red-wines over a seven-months period, following winemaking process season. A total of 2988 samples were retrieved from French wine cellars belonging to at least 12 wine regions and comprised red, white and rosé wines, as well as wine musts. Percentages of negative *B. bruxellensis* samples */* positive *B. bruxellensis* samples are indicated according to year, month, or week of analysis. Concentrations ranged between 0 and 1.58 x 10^7^ DNA copy. mL^-1^ (the highest concentration was reached on Week 39 in September 2023).

## 4. Conclusion

This study reported the development of a molecular tool for *B. bruxellensis* intact cells detection and quantification in the winemaking process and the first application of digital-PCR in this field to improve wine contaminants risk management. The method offers a rapid and robust quantification analysis for *B. bruxellensis* on which oenologists can rely to provide the appropriate treatments advice from grape harvest to bottling. Specificity of the analysis was improved in comparison to time-consuming conventional-plate culture, enabling to choose the appropriate treatment at the right moment to reduce chemical treatments and/or favor bioprotectants solutions. Our method has been proven successful for industrialization and analysis of a large number of naturally-contaminated samples, from different type of wines (red, white, rosé) and various matrix (grapes, wine must, fermenting wine, maturing wine). Our workflow is optimized to be representative of the total sample sent by wine producers, however, the sampling method in wine cellars must as well be representative and the complete sampling process needs to be considered. Overall, this work addresses a pressing need in food science to early detect microbial contaminants and offers a timely solution to enhance risk management, an emerging priority in a context of treatment reducing policies. This new digital-PCR method opens new perspectives for hygiene control of cellars by enabling detection of *B. bruxellensis* in wine as well as on surfaces (wipe sampling). The method could be extended to other microorganisms from the wine environment to improve our understanding of microbial dynamics in the wine and other beverage industry.

## CRediT authorship contribution statement

**Cécile Gruet:** Writing – original draft, Writing – review & editing, Methodology, Visualization, Conceptualization, Formal analysis. **Jeremy Di Mattia:** Writing – review & editing, Methodology, Visualization, Conceptualization, Data curation. **Magali Hiaumet:** Investigation, Methodology. **Dylan Pestel:** Investigation. **Caroline Araiz:** Writing – review & editing, Validation, Methodology. **Sarah Saadi:** Writing – review & editing, Investigation, Methodology. **Marie Ducousso**: Writing – review & editing, Supervision, Conceptualization. **Olivier Courot:** Writing – review & editing, Supervision, Conceptualization.

## Declaration of competing interest

The authors declare that they have no known competing financial interests or personal relationships that could have appeared to influence the work reported in this paper.

## Aknowledgments

We would like to thank Imane Youcef-Khodja for English proofreading and corrections, Benoît Desjardins for kind proofreading. This work made use of a strain collection provided by Microflora (affiliated with the Institut des Sciences de la Vigne et du Vin) and plate counting data were obtained thanks to the IGA analysis done by the Institut Coopératif du Vin.

## Notes

### Competing Interest Statement

The authors have declared no competing interest.

