## Supplementary Figures and Tables for "First application of digital-PCR in oenology for the specific detection of intact cells of *Brettanomyces bruxellensis* in the winemaking process"

*First co-authors

Pre-print

**Supplementary data**

**Figure S1. Preliminary extraction tests for *Brettanomyces bruxellensis* quantification by dPCR in wine.** (A) *B. bruxellensis* dPCR concentration after wine pellet washing with different solutions, compared to control sample with no washing. Tests were done in duplicates. The composition of the different solutions is not given as it remains a confidential information. (B) *B. bruxellensis* dPCR concentration with different washing solution C volumes. nW: non-washed, W200: 200 µL washing solution, W1500: 1.5 mL washing solution. (C) *B. bruxellensis* dPCR concentration depending on wine sampling volume, with or without pellet washing with 1.5 mL washing solution C. nW-100 : non-washed 100 µL of red wine, W-100 : washed 100 µL of red wine, W-500 : washed 500 µL of red wine, W-1000 : washed 1 mL µL of red wine, W-1500 : washed 1.5 mL of red wine, W-2000 : washed 2000 mL of red wine.

**Figure S2. Confirming experiment for pre-treatment protocol (Process B) and test for industrialization of the method.** (A) *B. bruxellensis* dPCR concentration according to the quantity of yeast added in 1,5 mL of wine depending on washing treatment. B: samples supplemented with 3/4 dilution of a *B. bruxellensis* liquid culture prepared at a known concentration, C: samples supplemented with 1/2 dilution, D: samples supplemented with 1/4 dilution, E: samples supplemented with 1/8 dilution, F: negative sample. Purple points correspond to washed samples according to the new protocol Process B (1.5 mL of washing solution C), gray points correspond to non-washed samples. Points and standard errors from conditions B, C, D and E correspond each to compiled data of three samples (each of them being tested in triplicates, giving n=6). Points and standard errors from condition F correspond to triplicates (n=3). $R^2$ = 0.98 for washed samples, $R^2$ = 0.97 for non-washed samples. (B) DNA extracts were the same samples than in (A), but they were analyzed on a 96-wells plate. *B. bruxellensis* dPCR concentration is indicated according to the quantity of yeast added in 1,5 mL of wine. $R^2$ = 0.99. (C) Correlation between results obtained with the 96-wells plate and the 24-wells plate. The correlation is statistically significant, Spearman correlation coefficient = 0.94.

**Table S1. *B. bruxellensis* dPCR concentration with or without treatment to detect only intact cells.** The process has been submitted for a FR patent (FR2401799) and is included into our sample pre-treatment before DNA extraction. 1.5 mL of wine were supplemented with different volume of *B. bruxellensis* pure gDNA at a known concentration. DNA extractions were performed after sample pre-treatment including or not the additional step for intact cells quantification. *B. bruxellensis* concentration into the dPCR reaction mix are indicated.

**Table S2. *Brettanomyces bruxellensis* quantification and yeast identity check in red wines according to two different methods.** A total of 42 other samples were analyzed. IGA method was developed by the Institut Coopératif du Vin and consists in conventional plate assays. dPCR analyses were made on DNA extract from 0.1mL of wine. Samples are sorted depending on CFU.mL-1 quantitites. For the IGA methods results were given in range rather than in absolute quantities. ND: Non detectable, meaning no *B. bruxellensis* was measured in the sample. Colonies on Petri dish were retrieved and analyzed by dPCR. "+" indicates presence of > 1 CFU on Petri dish or > 1 positive dPCR partition. "-" indicates no CFU on plates or no dPCR amplification.

Pre-print

**Figure S1**

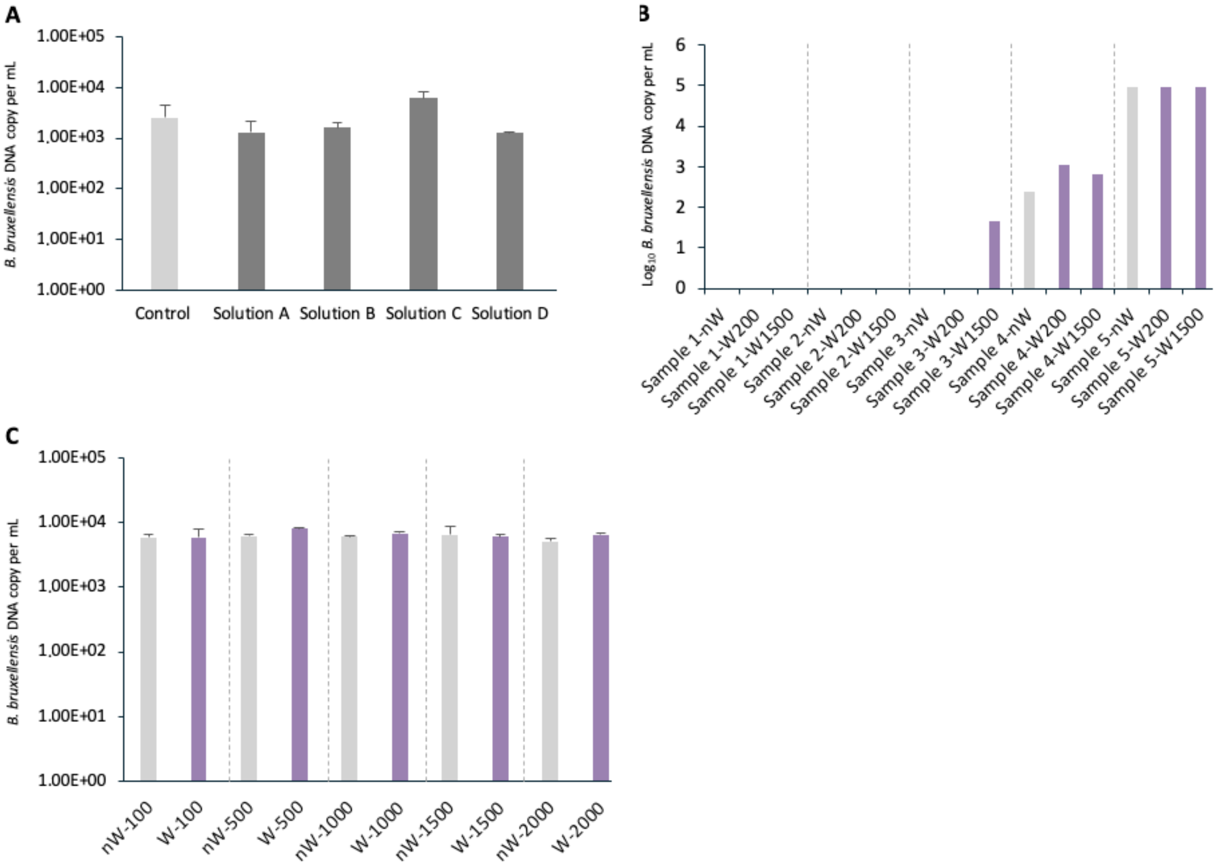


Pre-

**Figure S2**

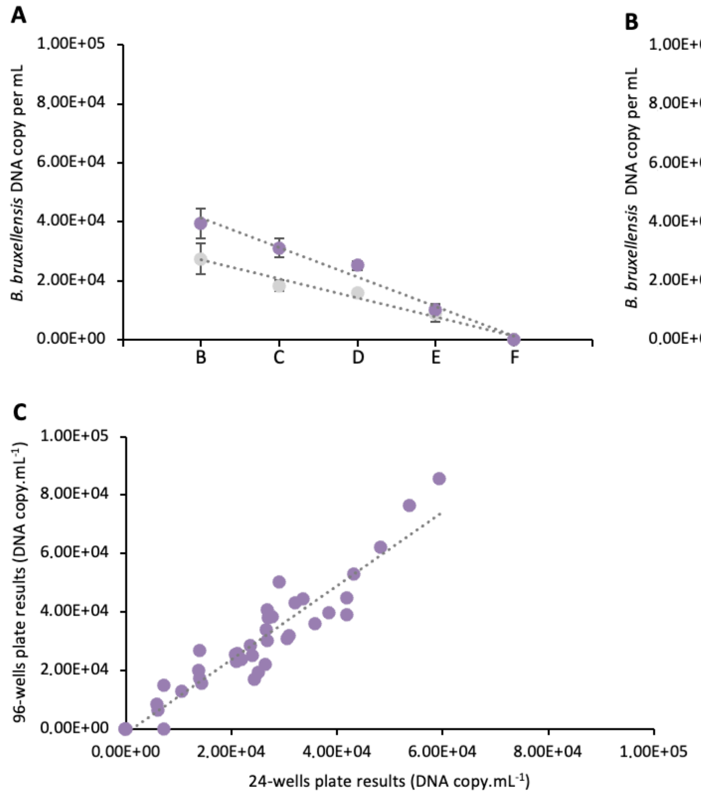


Pre-P

**Table S1.** ***B. bruxellensis*** **dPCR concentration with or without treatment to detect only intact cells.** The process has been submitted for a FR patent (FR2401799) and is included into our sample pre-treatment before DNA extraction. 1.5 mL of wine were supplemented with different volume of *B. bruxellensis* pure gDNA at a known concentration. DNA extractions were performed after sample pre-treatment including or not the additional step for intact cells quantification. *B. bruxellensis* concentration into the dPCR reaction mix are indicated.

| **Sample** | **Added volume of *B. bruxellensis* DNA (µL)** | ***B. bruxellensis* dPCR concentration (copy.µL-1)** | |
|---|---|---|---|
| | | Total DNA (without treatment) | Intact cells DNA (with treatment) |
| Wine 1 | 2 | 40.62 | 9.92 |
| | 2.5 | 50.69 | 22.86 |
| | 3 | 65.71 | 14.15 |

Pre-print

**Table S2.** ***Brettanomyces bruxellensis*** **quantification and yeast identity check in red wines according to two different methods.** A total of 42 other samples were analyzed. IGA method was developed by the Institut Coopératif du Vin and consists in a conventional plating method. dPCR analyses were made on DNA extract from 100 µL of wine. Samples are sorted depending on CFU.mL-1 quantitites. For the IGA methods results were given in range rather than in absolute quantities. ND: Non detectable, meaning no *B. bruxellensis* was measured in the sample. Colonies on Petri dish were retrieved and analyzed by dPCR. "+" indicates presence of > 1 CFU on Petri dish or > 1 positive dPCR partition. "-" indicates no CFU on plates or no dPCR amplification.

| Sample | IGA method ($CFU.mL^{-1}$) | Digital-PCR (DNA $copy.mL^{-1}$) | Colony dPCR |
|---|---|---|---|
| 1 | <1 | ND | - |
| 2 | <1 | ND | - |
| 3 | <1 | $7.40 \times 10^{1}$ | - |
| 4 | <1 | ND | - |
| 5 | <1 | ND | - |
| 6 | <1 | ND | - |
| 7 | <1 | $4.60 \times 10^{3}$ | + |
| 8 | <1 | $1.40 \times 10^{4}$ | + |
| 9 | <1 | ND | - |
| 10 | <1 | ND | - |
| 11 | <1 | ND | - |
| 12 | <1 | ND | - |
| 13 | <1 | ND | - |
| 14 | <1 | ND | - |
| 15 | <1 | ND | - |
| 16 | <1 | ND | - |
| 17 | <1 | ND | - |
| 18 | <1 | ND | - |
| 19 | <1 | ND | - |
| 20 | <1 | $7.24 \times 10^{3}$ | - |
| 21 | <1 | $8.32 \times 10^{3}$ | - |
| 22 | <1 | $3.85 \times 10^{3}$ | - |
| 23 | <1 | ND | - |
| 24 | <1 | ND | - |
| 25 | 10 - 100 | ND | - |
| 26 | 10 - 100 | ND | - |
| 27 | 10 - 100 | ND | - |
| 28 | 10 - 100 | ND | - |
| 29 | 10 - 100 | ND | - |
| 30 | 10 - 100 | ND | + |
| 31 | 10 - 100 | ND | - |
| 32 | 10 - 100 | ND | - |
| 33 | 100 - 1000 | $1.50 \times 10^{2}$ | + |
| 34 | 100 - 1000 | $1.00 \times 10^{4}$ | + |
| 35 | 100 - 1000 | $1.50 \times 10^{2}$ | + |
| 36 | 100 - 1000 | ND | - |
| 37 | 100 - 1000 | ND | - |

Pre-print

| | | | |
|---|---|---|---|
| 38 | 100 - 1000 | 2.30 x $10^3$ | + |
| 39 | 100 - 1000 | 5.40 x $10^2$ | + |
| 40 | 1000 | 1.12 x $10^3$ | + |
| 41 | 1000 | 1.10 x $10^3$ | + |
| 42 | 1000 | 1.50 x $10^5$ | + |

Pre-print
